# The differential distribution of bacteria between cancerous and noncancerous ovarian tissues in situ

**DOI:** 10.1101/562975

**Authors:** Qi Wang, Lanbo Zhao, Lu Han, Guoxing Fu, Xiaoqian Tuo, Sijia Ma, Qing Li, Yiran Wang, Dongxin Liang, Miaomiao Tang, Chao Sun, Qing Wang, Qing Song, Qiling Li

## Abstract

The female upper reproductive tract, including the uterus, fallopian tubes and ovaries, is believed to be a sterile environment. With the improvement of bacterial detection, the theory of the sterile female upper reproductive tract has been frequently challenged in recent years. However, thus far, no researchers have used ovaries as study targets. Six women who were diagnosed with ovarian cancer were included in the cancer group, and ten women who were diagnosed with a noncancerous ovarian condition (including three patients with uterine myoma and seven patients with uterine adenomyosis) were included in the control group. Immunohistochemistry staining using an antibacterial lipopolysaccharide (LPS) antibody was used to confirm the presence of bacteria in the ovarian tissues. In addition, 16S rRNA sequencing was used to compare the differences in the bacteria between ovarian cancer tissues and noncancerous ovarian tissues. BugBase and Phylogenetic Investigation of Communities by Reconstruction of Unobserved States (PICRUSt) were used to predict the functional composition of the bacteria. Bacterial LPS was present in ovarian cancer tissue and noncancerous ovarian tissue, which implied the presence of bacteria in ovarian tissue. When compared to the noncancerous ovarian bacteria at the phylum level, the cancerous ovarian bacteria were composed of increased Aquificae and Planctomycetes and decreased Crenarchaeota. When predicting metagenomes, gene functions associated with the potentially pathogenic and the oxidative stress-tolerant phenotype were enriched in the ovaries of the cancer group. Forty-six significantly different KEGG pathways existed in the ovarian bacteria of the cancer group compared to that of the control group. Different bacteria compositions were present in cancerous and noncancerous ovarian tissues.

**Author summary:** Abdominal solid viscera have always been believed to be absolutely sterile. With the improvement of bacterial detection, this concept is being challenged. Researchers found some bacteria existed in endometrial diseases, the question of whether the ovaries are sterile is still unclear. Therefore, we assume that the bacteria in ovarian tissue are associated with ovarian cancer. When compared to the noncancerous ovarian bacteria at the phylum level, the cancerous ovarian bacteria were composed of increased Aquificae and Planctomycetes and decreased Crenarchaeota. When predicting metagenomes, gene functions associated with the potentially pathogenic and the oxidative stress-tolerant phenotype were enriched in the ovaries of the cancer group. Forty-six significantly different KEGG pathways existed in the ovarian bacteria of the cancer group compared to that of the control group. Different bacteria compositions were present in cancerous and noncancerous ovarian tissues.

## Introduction

Abdominal solid viscera, including the pancreas, kidney, spleen, liver and ovary, have always been believed to be absolutely sterile. However, this concept is being challenged. Leore *et al*. found that the bacteria in pancreatic tumors could mediate tumor resistance to the chemotherapeutic drug gemcitabine(1). S. Manfredo Vieira *et al*. confirmed that *Enterococcus gallinarum* can translocate to the lymph nodes, liver and spleen and drive autoimmunity(2).

The upper female reproductive tract, including the uterus, fallopian tubes and ovaries, has been believed to be absolutely sterile due to the obstacle of the cervix, which is also being challenged. The change in mucins in the cervix during the menstrual cycle may lead to the passage of bacteria(3, 4). In addition, research has confirmed that the uterus and fallopian tubes represent a functionally united peristaltic pump under the endocrine control of the ovaries(5), which may aid the bacteria to enter the endometrium, fallopian tubes, and ovaries.

With the improvement of bacterial detection, researchers have been investigating the upper reproductive tract. Verstraelen *et al*. aimed to explore the presence of a uterine bacteria using a barcoded Illumina paired-end sequencing method targeting the V1-2 hypervariable region of the 16S RNA gene(6). Fang *et al*. revealed diverse intrauterine bacterias in patients with endometrial polyps using barcoded sequencing (7). Miles and Chen also investigated the bacteria of the reproductive tract in women undergoing hysterectomy and salpingo-oophorectomy using the 16S RNA gene(4, 8). However, all of the abovementioned researchers used endometrial diseases as their research targets, so the question of whether the ovaries are sterile is still unclear.

In recent years, the bacteria of tumor tissues have become a hot topic for researchers. Aleksandar *et al*. confirmed that Fusobacterium was enriched in colorectal tumors (9). In addition, Bullman *et al*. discovered that the colonization of human colorectal cancers with Fusobacterium is maintained in distal metastases and bacteria stability between paired primary and metastatic tumors(10). Therefore, we assume that the bacteria in ovarian tissue are associated with ovarian cancer.

In this study, we used immunohistochemistry staining and 16S rRNA sequencing to confirm the presence of bacteria in the ovaries. First, we compared the differences in the ovarian bacteria and its predicted function between cancerous and noncancerous ovarian tissues.

## Results

### Participant patients

Sixteen patients who were undergoing oophorectomy or hysterectomy and salpingo-oophorectomy were included in this study. In this study, ten women who were diagnosed with benign endometrial conditions with noncancerous ovaries (including three patients with uterine myoma and seven patients with uterine adenomyosis) were set as the control group, and six women who were diagnosed with ovarian cancer (including two patients who were diagnosed in stage II and four patients who were diagnosed in stage III) were set as the cancer group. All diagnoses were based on final surgical pathology after oophorectomy or hysterectomy and salpingo-oophorectomy. Compared with the control group, the age, menopausal status, parity, history of hypertension and history of diabetes in patients diagnosed with ovarian cancer were not significantly different (Table 1).

**Table 1.**
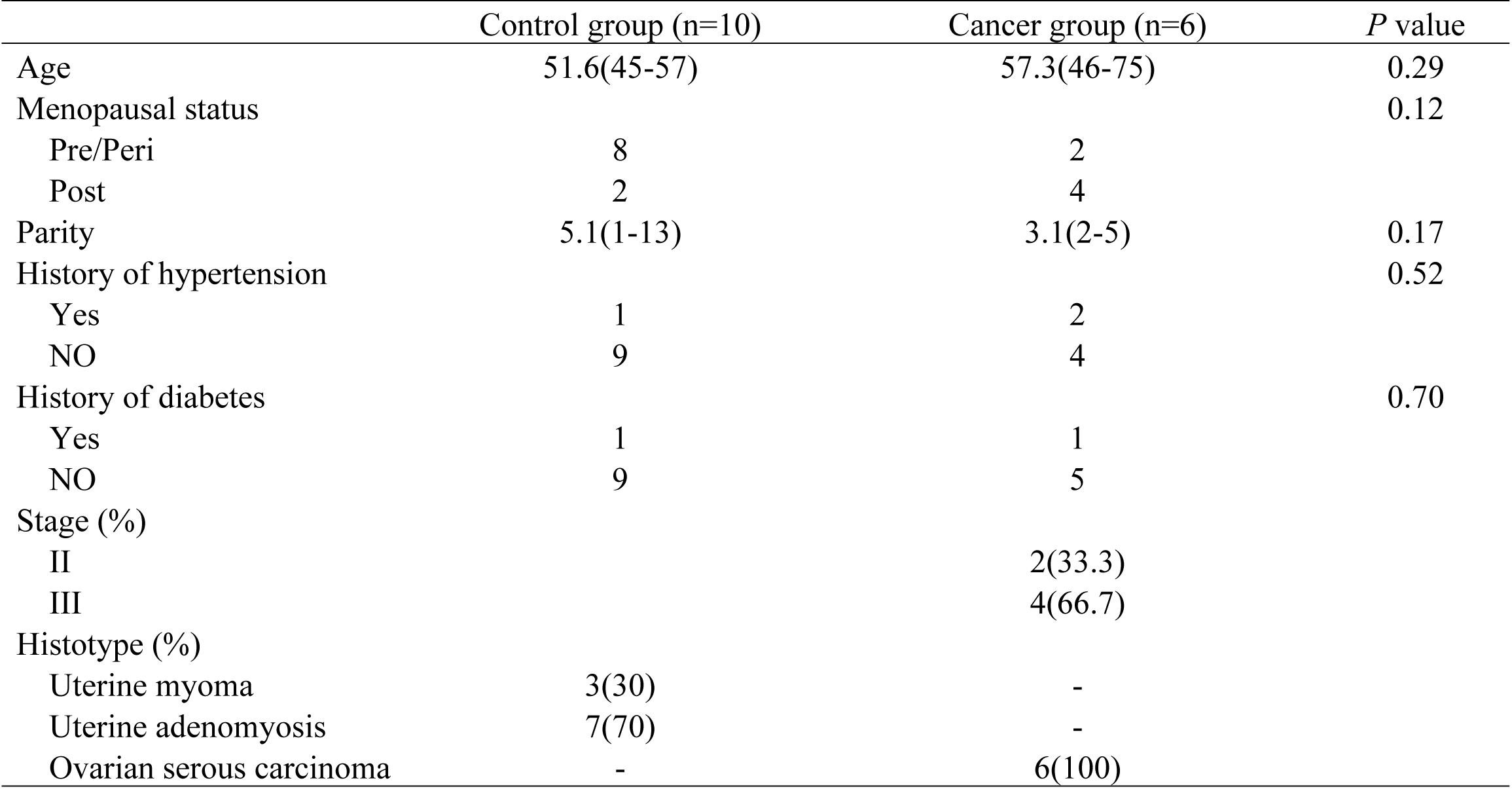
Clinical characteristics of patients enrolled in the study.

|  | Control group (n=10) | Cancer group (n=6) | *P* value |
| --- | --- | --- | --- |
| Age | 51.6(45-57) | 57.3(46-75) | 0.29 |
| Menopausal status |  |  | 0.12 |
| Pre/Peri | 8 | 2 |  |
| Post | 2 | 4 |  |
| Parity | 5.1(1-13) | 3.1(2-5) | 0.17 |
| History of hypertension |  |  | 0.52 |
| Yes | 1 | 2 |  |
| NO | 9 | 4 |  |
| History of diabetes |  |  | 0.70 |
| Yes | 1 | 1 |  |
| NO | 9 | 5 |  |
| Stage (%) |  |  |  |
| II |  | 2(33.3) |  |
| III |  | 4(66.7) |  |
| Histotype (%) |  |  |  |
| Uterine myoma | 3(30) | - |  |
| Uterine adenomyosis | 7(70) | - |  |
| Ovarian serous carcinoma | - | 6(100) |  |

### The presence of bacteria in the ovaries

To confirm the presence of bacteria in ovaries using non-PCR-based methods, we performed immunohistochemistry staining using an antibacterial LPS antibody. The results showed that bacterial LPS were present in the cancerous ovarian tissue and noncancerous ovarian tissue, which implied the presence of bacteria in ovarian tissue (Fig. 1).

**Fig 1.**
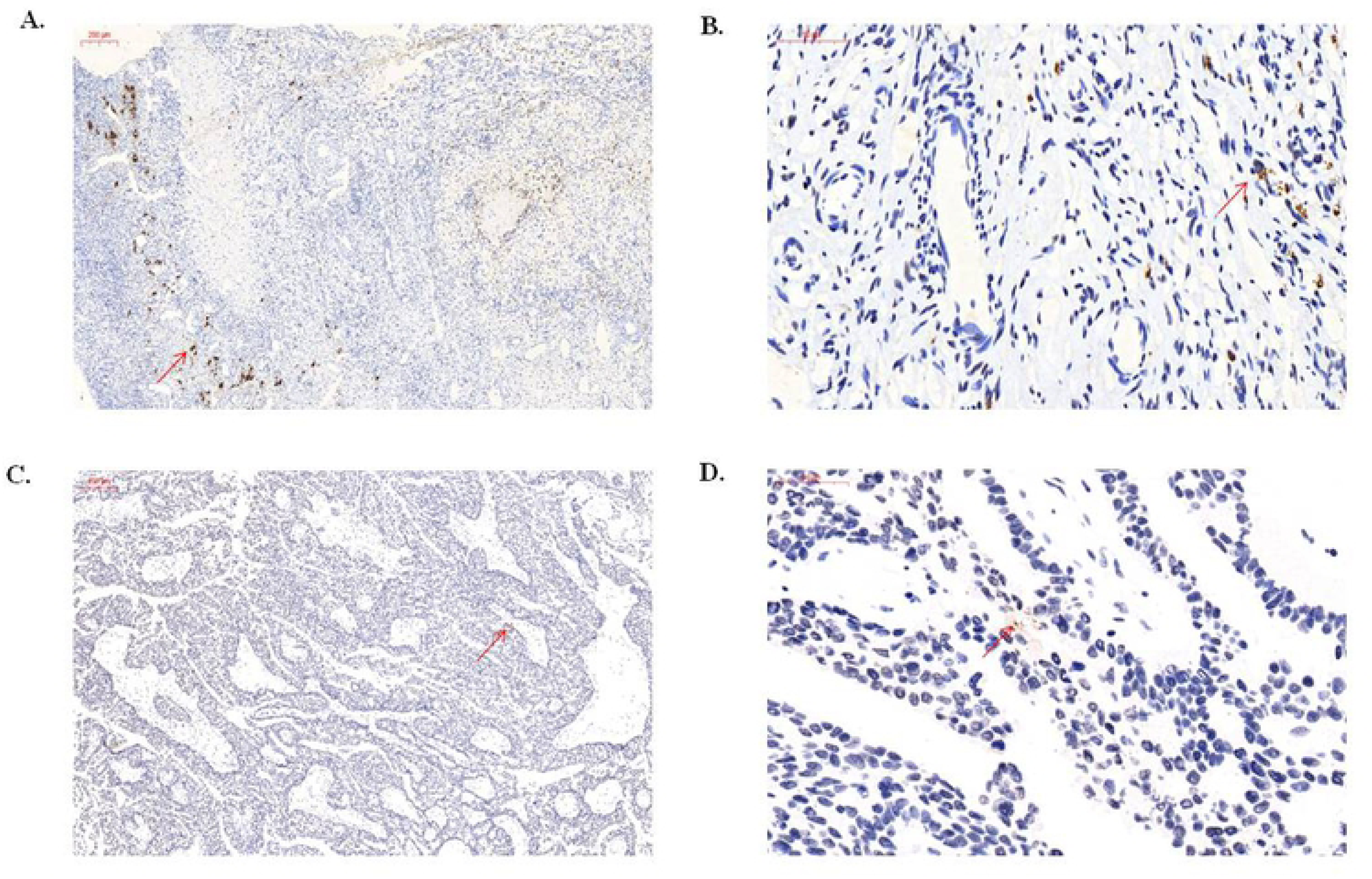
Immunohistochemistry of ovaries using an antibacterial LPS antibody. A. control group (10x). Scale bars, 200 µm. B. control group (40x). Scale bars, 50 µm. C. cancer group (10x). Scale bars, 200 µm. D. cancer group (40x). Scale bars, 50 µm. Arrows point to LPS staining in the ovarian tissue.

### Ovarian bacterial richness and diversity between the cancer and control groups

To detect the ovarian bacterial species richness and diversity between the two groups, we analyzed the alpha diversity of the microbes. The observed number of species in the ovarian cancer tissues was lower than that in the ovaries of the control group, but the difference was not significant. Moreover, we found that not only the bacterial species richness (represented by the Chao 1 index and the ACE index) but also the diversity (represented by the Shannon Index, the Simpson Index and the Evenness Index) in the ovarian cancer group were not significantly different from those in the control group (Fig. 2).

**Fig 2.**
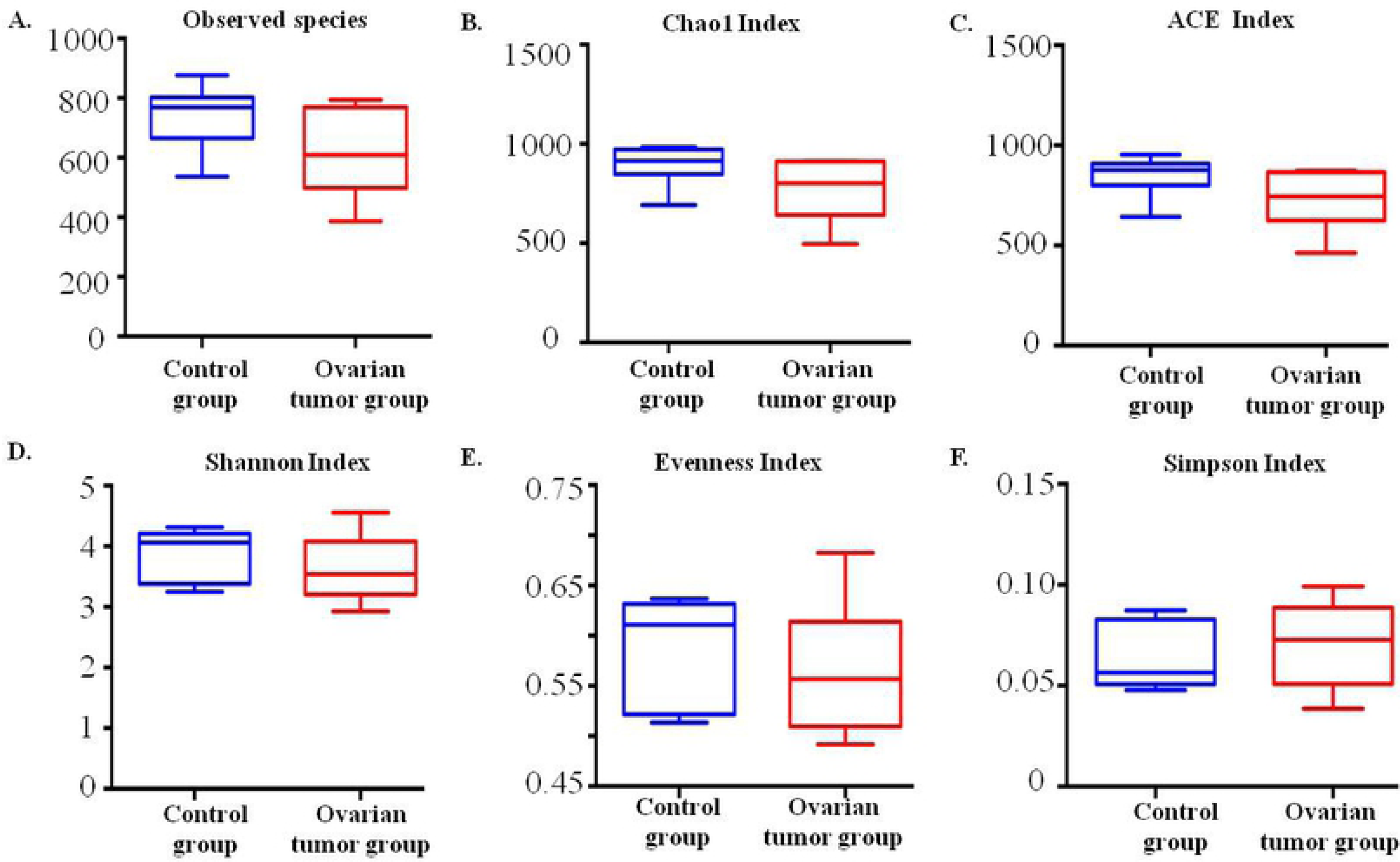
Bacterial richness and diversity in the cancer and control groups revealed by 16S rRNA sequencing. A. Observed species index P=0.06; B. Chao 1 index P=0.06; C. ACE index P=0.06; D. Shannon index P=0.32; E. Evenness index P=0.48; F. Simpson index P=0.46.

### Ovarian bacteria characterization between the cancer and control groups

To understand the ovarian bacteria in the cancer and control groups, we performed deep sequencing of the V3-V4 16S rRNA region of all sixteen collected samples. In the ovaries, our results showed that Proteobacteria was the most abundant phylum (67.1% in the control group and 67.20% in the cancer group). Firmicutes was the second most abundant phylum (23.77% in the control group and 23.82% in the cancer group), and the third most abundant phylum was Bacteroidetes (3.26% in the control group and 3.41% in the cancer group) (Fig. 2A, 2B). At the species level, the ovarian bacterial communities were dominated by *Halobacteroides halobius* (14.53%), followed by *Gemmata obscuriglobus* (11.07%) and *Methyloprofundus sedimenti* (10.69%) in the control group. The ovarian bacterial communities in the cancer group were dominated by *Gemmata obscuriglobus* (13.89%), followed by *Halobacteroides halobius* (11.99%) and *Methyloprofundus sedimenti* (11.12%) (Fig. 3).

**Fig 3.**
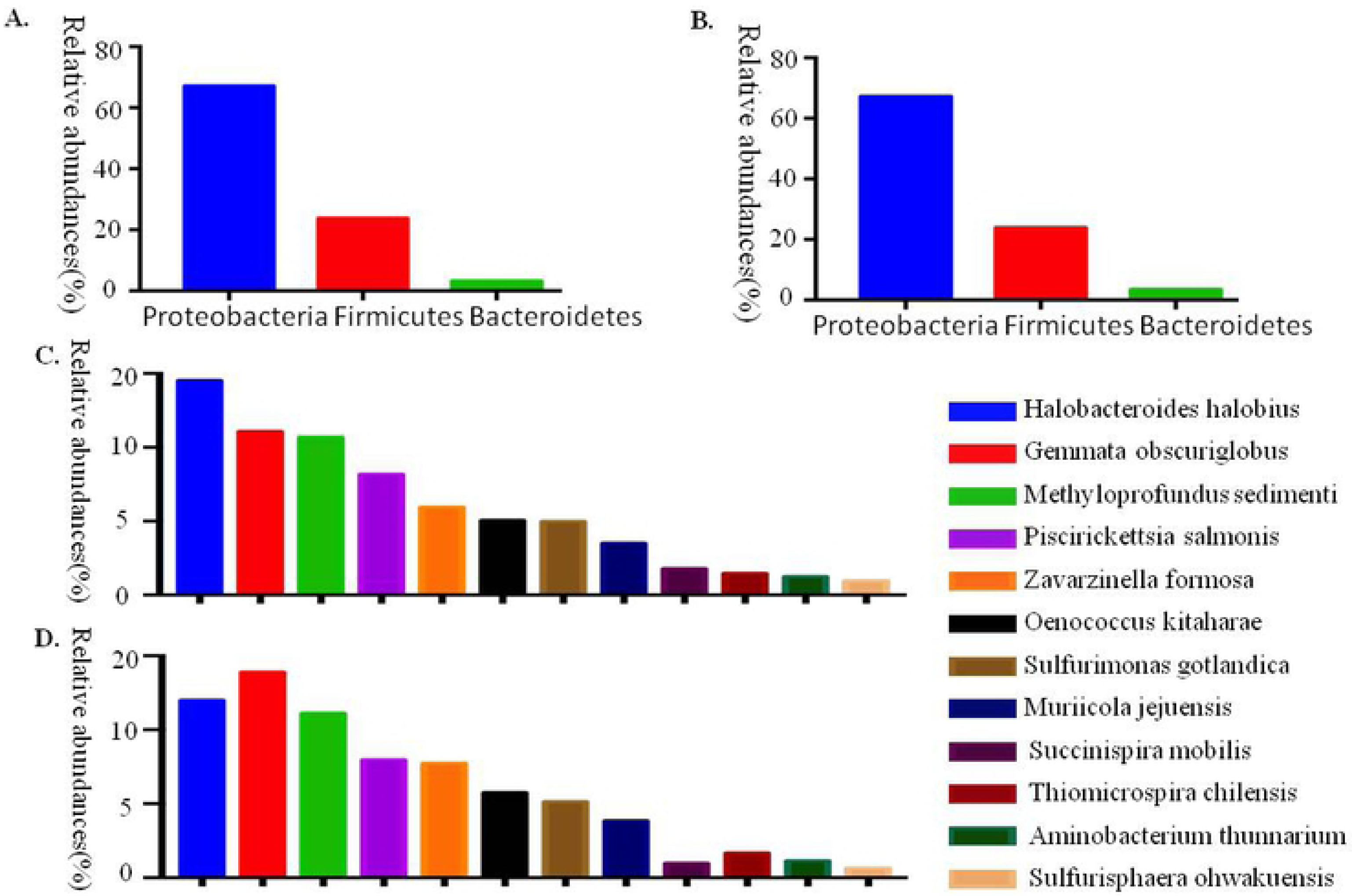
The relative abundance of phyla (>1%) and the 12 most abundant bacterial species in the ovarian samples. A. The relative abundance of the phyla (>1%) in the ovaries of the patients in the control group. B. The relative abundance of the phyla (>1%) in the ovaries of patients with ovarian cancer. C. The relative abundances of the 12 most abundant bacterial species in the ovaries of the control patients. D. The relative abundances of the 12 most abundant bacterial species in the ovaries of ovarian cancer patients.

### Ovarian bacterial community composition differences between the cancer and control groups

We carried out a comparison of differences in the overall bacterial communities using PCoA, which showed that the ovarian bacteria of the control group displayed some differences compared to that of the cancer group (Fig. 4A and 4B).

**Fig 4.**
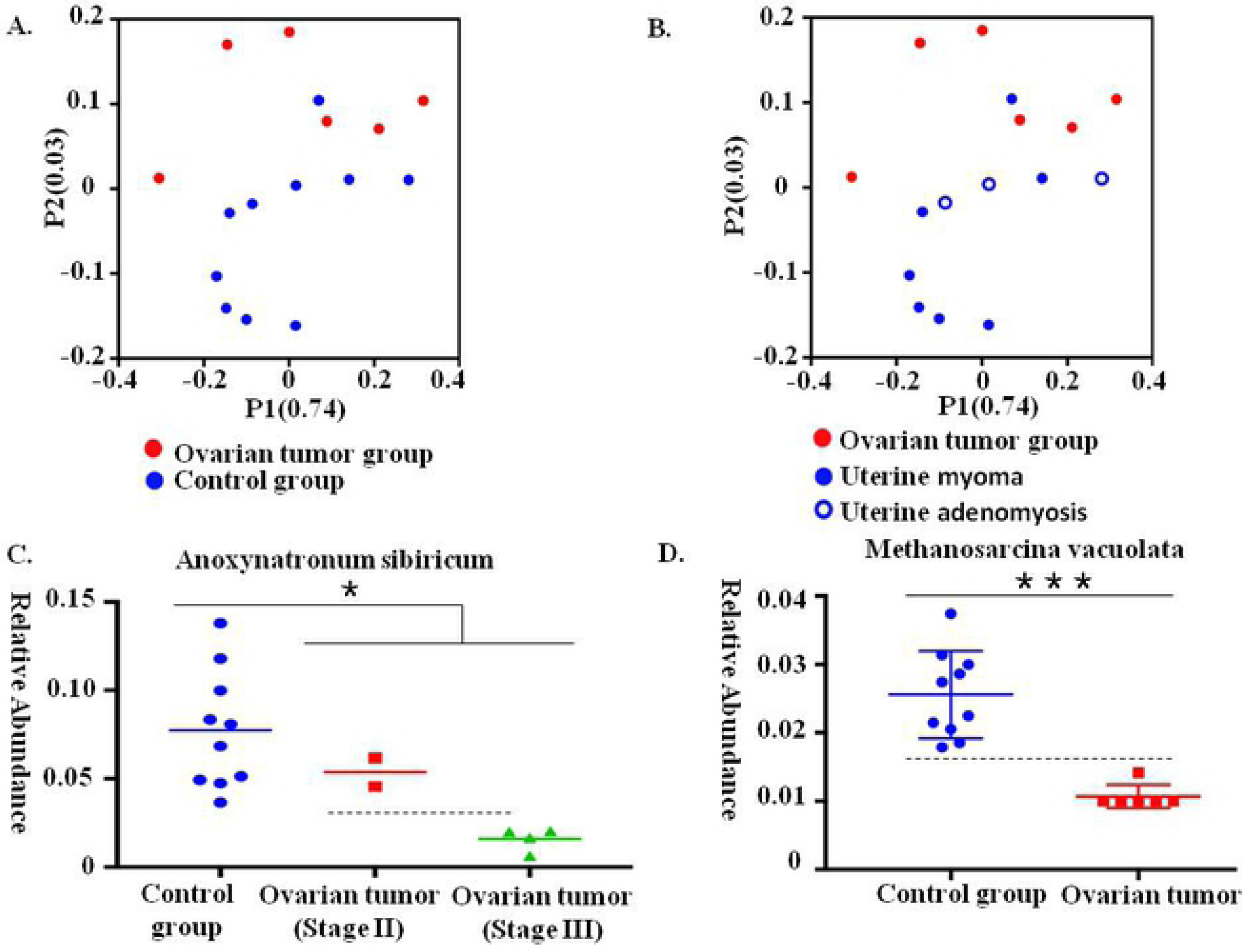
Communities clustered using PCoA and the relative abundance of *Anoxynatronum sibiricum* and *Methanosarcina vacuolata*. A. Communities were clustered using PCoA. PC1 and PC2 are plotted on the x and y axes. The red block is equal to a sample in the ovarian cancer group. The blue circle is equal to a sample in the control group. The samples from the ovarian cancer group can be separated from other samples in the control group. B. Communities clustered using Principal Component Analysis (PCoA). PC1 and PC2 are plotted on the x and y axes. The red block is equal to a sample in the ovarian cancer group. The blue solid circle is equal to a sample from a patient with uterine myoma, and the blue hollow circle is equal to a sample of a patient with uterine adenomyosis. C. The relative abundance of *Anoxynatronum sibiricum*. D. The relative abundance of *Methanosarcina vacuolata*.

### Ovarian bacterial composition at different levels in the cancer and control groups

To detect the differences in ovarian bacteria between the seventeen samples, we analyzed the ovarian bacterial composition at different levels in the cancer and control groups. At the phylum level, the relative abundance of Aquificae and Planctomycetes in the cancer group was higher than that in the control group (*P*=0.017, *P*= 0.023, respectively), and the abundance of Crenarchaeota in the cancer group was lower than that in the control group (*P*=0.023). At the class level, the relative abundance of Spartobacteria sequences was significantly higher in the ovarian communities of the cancer group (*P*=0.026), whereas that of Sphingobacteriia was significantly lower (*P*=0.039). At the genus level, we found that the relative abundance of Pelagicoccus, Haloferula, Volucribacter, Blastococcus and Defluviitoga in the ovarian communities of the cancer group was significantly lower than that of the control group, and the relative abundance of Zavarzinella, Photorhabdus and Mesotoga in the ovarian communities of the control group was significantly lower than that of the cancer group. At the species level, the relative abundance of *Luteolibacter cuticulihirudinis*, *Aureimonas phyllosphaerae*, *Azonexus hydrophilus*, *Anaerostipes rhamnosivorans*, *Calditerricola yamamurae*, *Peptoniphilus methioninivorax*, *Vulcanisaeta thermophile*, a subspecies of *Staphylococcus capitis*, *Mycoplasma genitalium*, *Sulfurospirillum halorespirans*, *Methylomicrobium album*, Caldicoprobacteroshimai, *Thermogemmatispora foliorum*, *Mycoplasma equigenitalium*, *Bifidobacterium subtile* and *Rhodopirellula rubra* was significantly higher in the ovarian communities of the ovarian cancer group, whereas the relative abundance of thirty-seven other species was lower than that in the control group (Table 2). In particular, the relative abundance of *Anoxynatronum sibiricum* may be associated with the stage of the tumor (Fig. 4C), and *Methanosarcina vacuolata* may be used to diagnose ovarian cancer (Fig. 4D).

**Table 2.**
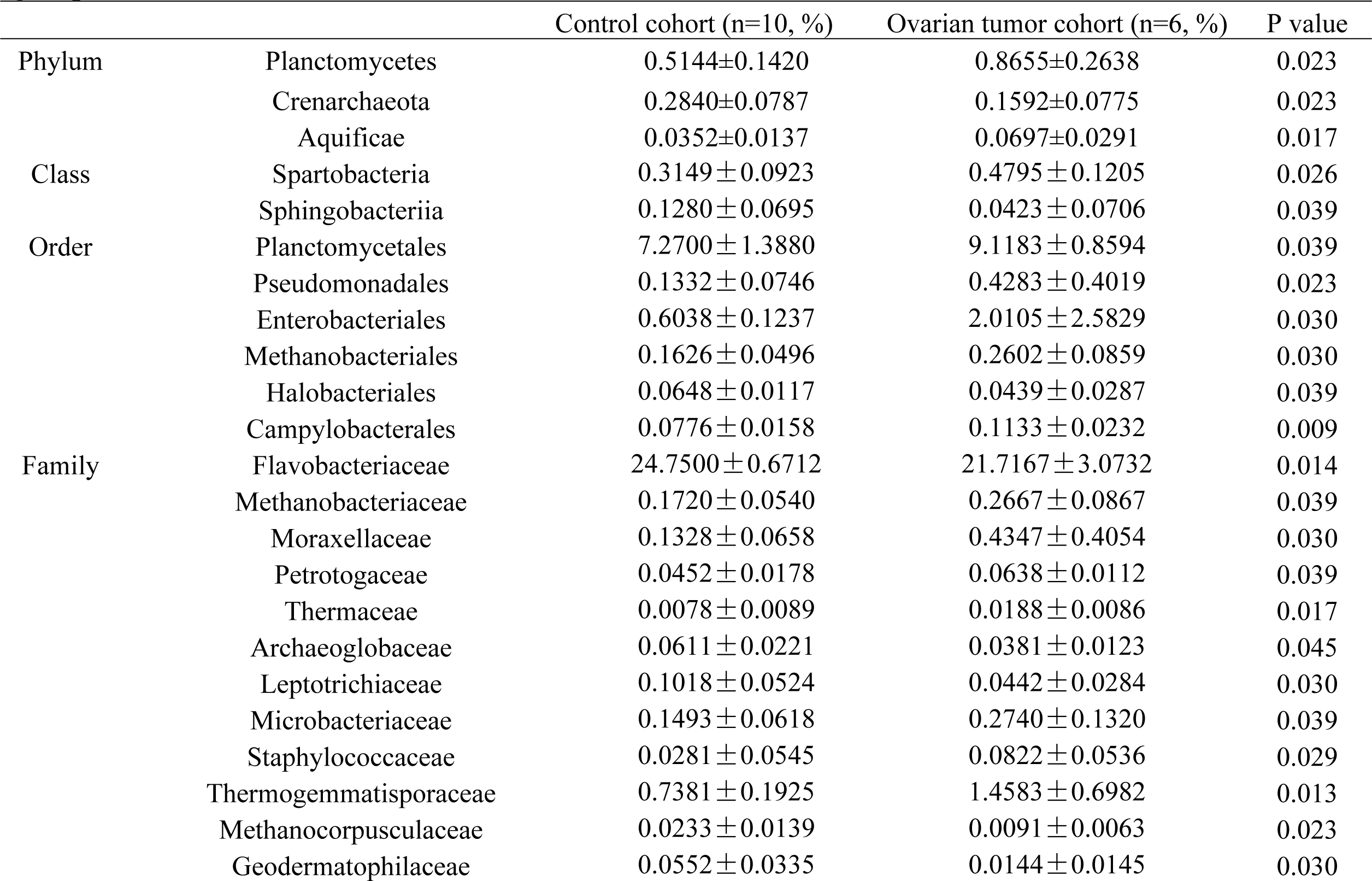
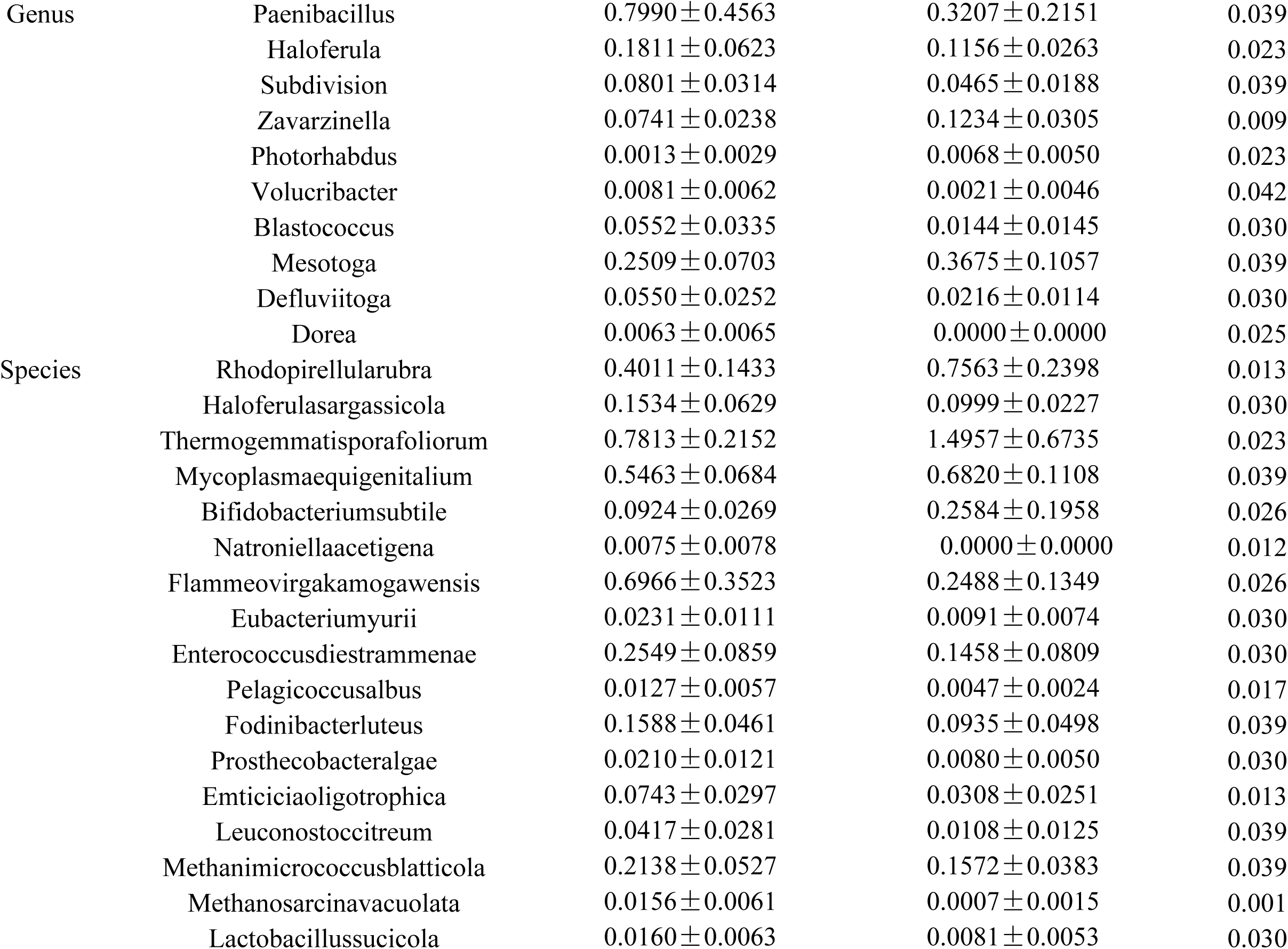
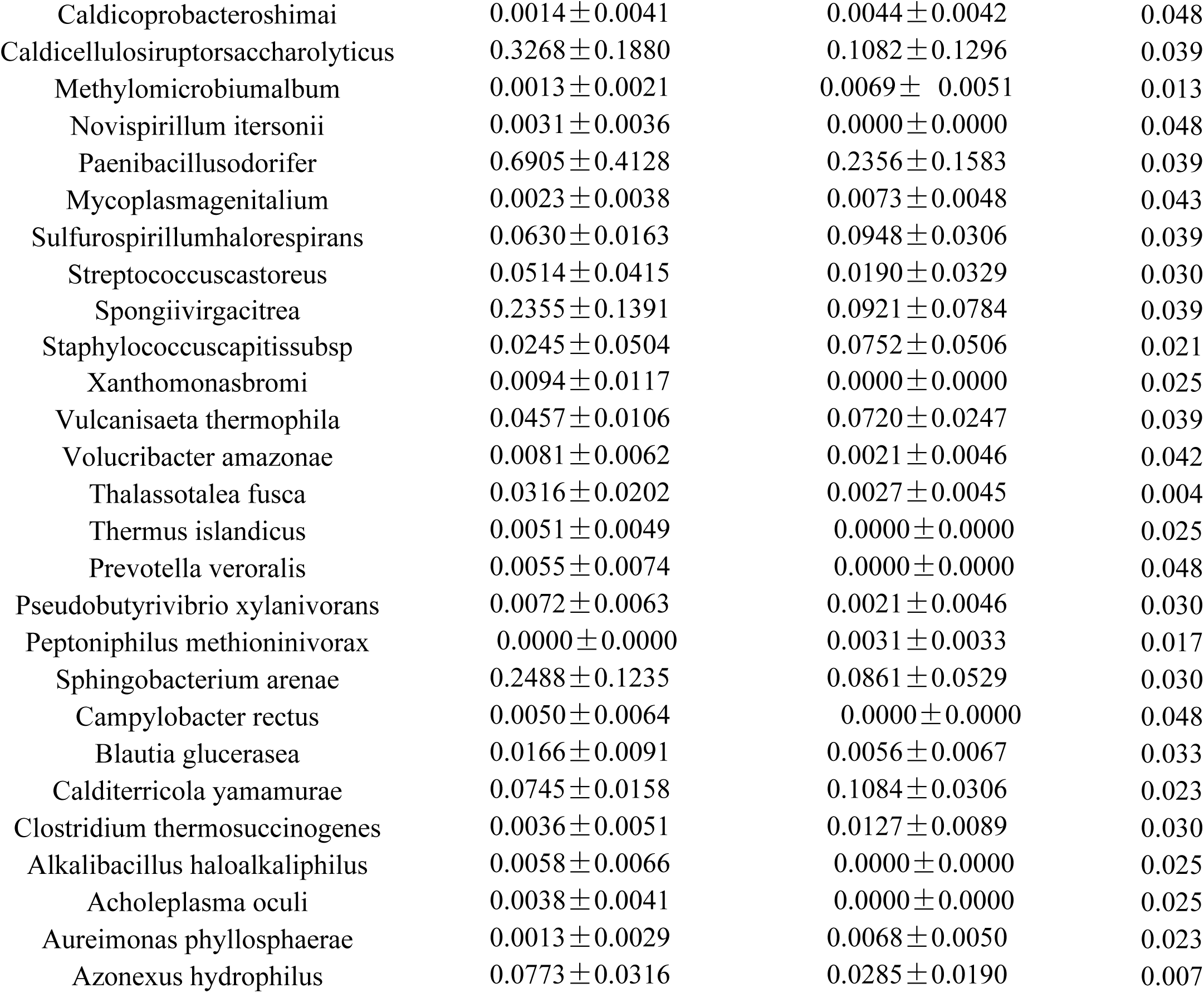
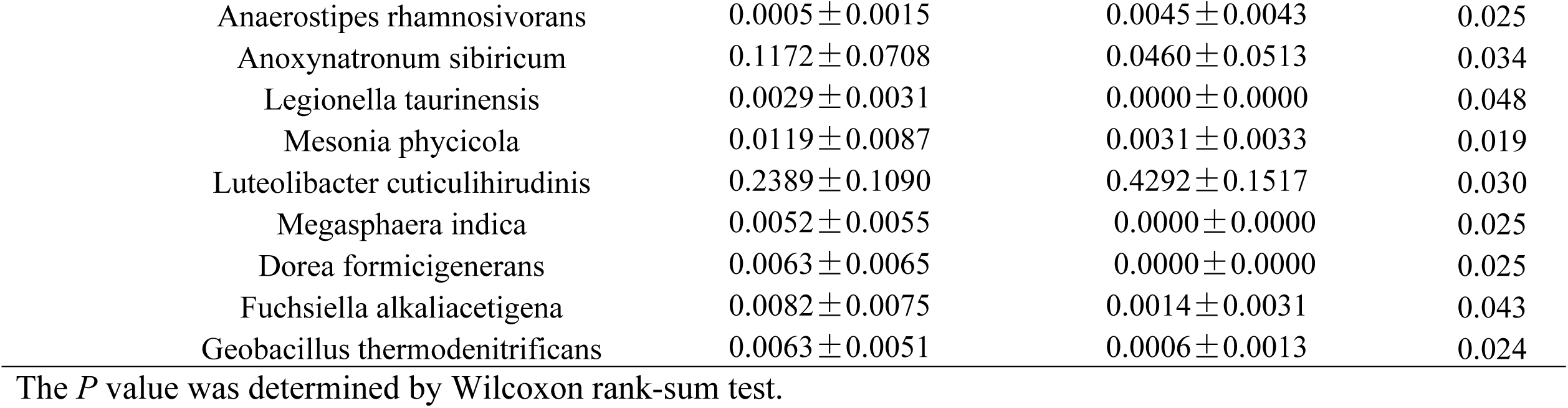
Differential relative abundance of the taxa in ovarian communities between patients in cancer and control group.

### Predicted function of the ovarian bacteria shows phenotypic conservation between the cancer and control groups

BugBase identified that gene functions associated with the potentially pathogenic and the oxidative stress-tolerant phenotype were enriched in the ovaries of the cancer group (Wilcoxon signed-rank test, *P* = 0.02 and *P* = 0.002). The aerobic, anaerobic, facultative anaerobic, gram-positive, and gram-negative phenotypes; mobile elements; and biofilm formation of the ovarian bacteria showed no significant difference between the ovarian cancer and control groups (Fig. 5). PICRUSt was used to identify the KEGG pathways between the bacteria of ovaries in the cancer and control groups and found 46 different KEGG pathways. The ovaries in the cancer group showed increased pathways related to streptomycin biosynthesis, carbon fixation in photosynthetic organisms, glycosphingolipid biosynthesis-globo series, cyanoamino acid metabolism, glycerophospholipid metabolism, butirosin and neomycin biosynthesis, other glycan degradation, biosynthesis of vancomycin group antibiotics, polyketide sugar unit biosynthesis, the pentose phosphate pathway, transporters, tuberculosis, starch and sucrose metabolism, fructose and mannose metabolism, phenylalanine metabolism, lysosomes, glycosaminoglycan degradation, pentose and glucuronate interconversions, pyruvate metabolism, amino sugar and nucleotide sugar metabolism, galactose metabolism, biosynthesis of ansamycins, methane metabolism, membrane and intracellular structural molecules, metabolism of cofactors and vitamins, glutamatergic synapse, and the cell cycle. However, the bacteria in ovarian cancer tissue showed reduced alpha-linolenic acid metabolism, biosynthesis of unsaturated fatty acids, bacterial secretion system, proximal tubule bicarbonate reclamation, prion diseases, secretion system, carbon fixation pathways in prokaryotes, unknown functions, other ion-coupled transporters, sulfur metabolism, biotin metabolism, protein kinases, ubiquinone and other terpenoid-quinone biosynthesis, two-component system, folate biosynthesis, cell motility and secretion, citrate cycle (TCA cycle) and ribosome biogenesis in eukaryotes (Fig. 6).

**Fig 5.**
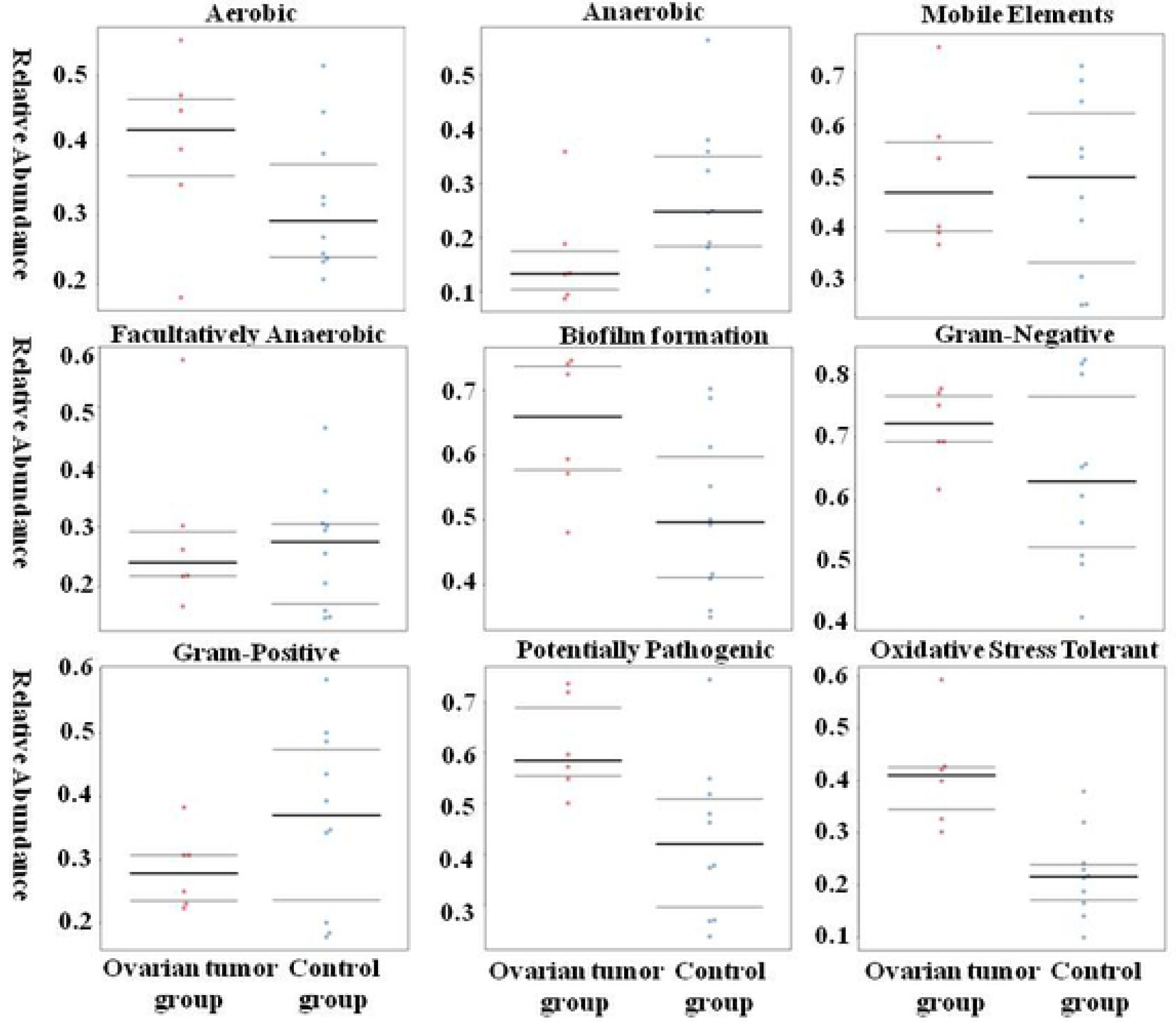
BugBase analysis of predicted metagenomes. The potentially pathogenic and oxidative stress-tolerant phenotype of the ovaries in the cancer group was stronger than that of the control group. (Wilcoxon signed-rank test, *P* = 0.02 and *P* = 0.002).

**Fig 6.**
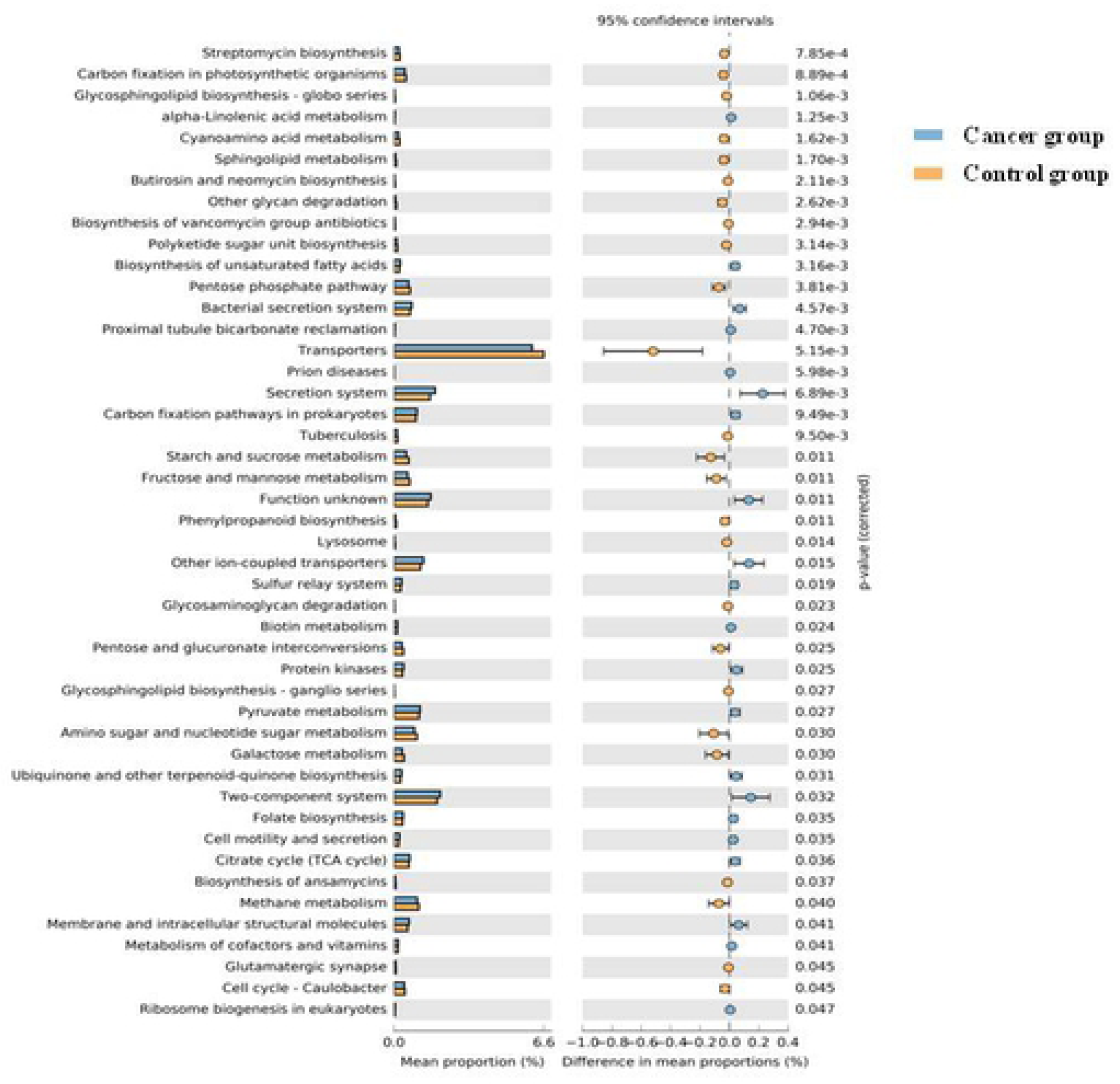
The significantly different KEGG pathways between the cancer and control groups by PICRUSt analysis.

## Discussion

Ovarian cancer (OC) is the seventh most commonly diagnosed cancer among women that could affect fertility(11). Most ovarian cancer patients are diagnosed at stages III and IV, and the 5-year survival rate is less than 30% (12). Researchers have confirmed that the abdominal solid viscera, including the liver, pancreas and spleen, are not absolutely sterile, and the bacteria exists in the upper female reproductive tract as a result of leakage from the cervix(1, 4). However, the ovaries are still not a research target. The question remains unanswered that whether the ovaries, as one of the abdominal solid viscera, have a bacteria and whether the bacteria has an association with ovarian cancer.

In this study, we first confirmed the presence of bacteria in the ovaries. In addition, we detected significant differences in the ovarian bacteria of patients with ovarian cancer when compared with samples from noncancerous women. At the genus level, we found a lower relative abundance of Pelagicoccus, Haloferula, Volucribacter, Blastococcus and Defluviitoga in ovarian cancer and a higher relative abundance of Zavarzinella, Photorhabdus and Mesotoga in noncancerous ovaries. The genus Paenibacillus was previously confirmed to be present in the human body, including in the blood, ascetic fluid, cornea pericardium, and cerebrospinal fluid(13). The genus Photorhabdus has been related to soft tissue infection and bacteremia(14). The genus Dorea exists in human gut mucosa and tumor tissue and is related to colorectal cancer(15, 16). Another genus was first found present in the human body and had a potential association with ovarian cancer. We predicted ovarian metagenomes and found that the pathogenic and oxidative stress-tolerant phenotype was enriched in the ovaries of the cancer group. Moreover, the enhanced function of cell growth and death in the KEGG pathway was detected in ovarian cancer, which could explain how the bacteria of the ovaries influenced the occurrence and development of the tumor.

To avoid bacterial contamination, all instruments used were sterilized, and the reagent we used was new. When operating, the surgeon wore an autoclaved mask, cap and suit and did not talk. The sample did not touch anything in the operating room except for the tweezers and was immediately put into the sterilized tube. When the sample was transferred to the laboratory, as many of the procedures as possible were performed on the asepsis work table except the procedures that required large equipment, such as centrifugal machines and sequencers. More importantly, we used ovaries from patients with benign uterine disease as the control group to counteract possible contamination.

There are three possible reasons to explain the origination of the ovarian bacteria. First, a new opinion is that the upper female reproductive tract is not sterile(4), and different bacteria exist throughout the female reproductive tract, forming a continuum from the vagina to the ovaries(12). The bacteria in the ovaries may originate from the fallopian tubes, uterine cavity, cervix canal or vagina, which is in contact with the outside environment. Second, the bacteria in the upper female reproductive tract, including the ovaries, may be endosymbiotic and separated from other bacteria and the outside environment(4). Third, the blood and abdominal cavity may be the potential source of the ovarian bacteria, which is our hypothesis.

In this study, we found the presence of bacteria in the ovaries and differences in the ovarian bacteria between patients with ovarian cancer and noncancerous women, which raises further questions that need to be solved. Where did the bacteria originate from? What is the association between the bacteria in the ovaries, uterus, fallopian tubes, vagina, and the outside environment? Are the ovarian bacteria always present? The ovaries are connected and open to the abdominal cavity; did the bacteria transfer from the abdominal cavity and the surface of the organs? Moreover, another doubt is whether the ovarian bacteria is associated with ovulation, ovarian failure, ovarian cysts, polycystic ovarian syndrome and so on. Do the ovarian bacteria drive the occurrence of ovarian cancer or does ovarian cancer change the ovarian bacteria? All of the above questions point to the direction of our future research.

There are some limitations to our study. The first limitation is that we could not collect the ovaries from healthy patients for ethical reasons. Therefore, we used the noncancerous ovaries from patients with benign uterine disease (including uterine myoma and adenomyosis) as the control group. Another limitation of this study is the small sample size, which may limit further analysis and influence the accuracy of the results. However, it is the preliminary study to detect the ovarian bacteria in patients with ovarian cancer, and we will conduct further explorations with larger sample sizes.

The ovaries contained several kinds of bacteria and were not sterile in a noninflammatory environment. There were significant differences between the ovarian bacterial compositions of patients in the cancer and control groups.

## Materials and Methods

### Ethics Statement

This study protocol was approved by the Ethics Committee of the First Affiliated Hospital of Xi’an Jiaotong University of China (Approval number: XJTU1AF2018LSK-139). The written informed consents were obtained from all patients participated in the study. All human subjects were adult.

### Patient characteristics

Sixteen patients were enrolled at the First Affiliated Hospital of Xi’an Jiaotong University. Patients with any of the following criteria were included in our study: patients undergoing oophorectomy by standard surgical approach and patients undergoing hysterectomy and salpingo-oophorectomy for benign uterine disease (including uterine myoma and uterine adenomyosis) or any stage of ovarian cancer. The exclusion criteria were as follows: patients who were pregnant or nursing, patients who took antibiotics within two months before surgery, and patients who had a fever or elevated inflammatory markers.

### Sample collection

Once removed, the ovaries were cut into approximately 1-cm thick ovarian tissue samples using a pair of sterile new tweezers without touching anything else. Then, the collected sample was placed into a sterile tube and placed in liquid nitrogen. Specimens were then transferred to the laboratory and stored at –80°C.

### Immunohistochemistry for bacterial lipopolysaccharide (LPS) in ovaries

Immunohistochemistry staining was performed on 5 μm serial sections from routine formalin-fixed, paraffin-embedded (FFPE) tissues. The samples were deparaffinized and rehydrated, and antigen retrieval was performed by microwave treatment for 10 minutes in EDTA buffer (pH 9.0). Endogenous peroxidase activity was stopped by incubating samples with 0.3% hydrogen peroxide in PBS for 20 minutes. A DAB substrate kit was used to detect HRP (Zytomed Systems, Berlin, Germany). A ZytoChem Plus HRP Polymer Anti-Rabbit secondary antibody was used according to the manufacturer’s instructions (Zytomed Systems). To find the bacteria, the antibody to LPS core (Hycult Biotech, Uden, Netherlands; Clone WN1 222-5) was used at a concentration of 1:300 overnight at 4°C(1).

### 16S rRNA sequencing

DNA extractions were performed by using the Mag-Bind® Stool DNA 96 Kit (Omega Biotek, Norcross, USA). DNA was quantified using the QuantiFluor dsDNA System (Promega, Madison, USA). The libraries were prepared using an Illumina 16S Metagenomic Sequencing kit (Illumina, Inc., San Diego, USA) according to the manufacturer’s protocol. The V3-V4 region of the bacterial 16S rRNA gene sequences was amplified using the primer pair containing the gene-specific sequences and Illumina adapter overhang nucleotide sequences. The full-length primer sequences were as follows: 16S Amplicon PCR Forward primer: 5’ TCGTCGGCAGCGTCAGATGTGTATAAGAGACAG-[CCTACGGGNGGCWGCAG] and 16S Amplicon PCR Reverse primer: 5’ GTCTCGTGGGCTCGGAGATGTGTATAAGAGACAG-[GACTACHVGGGTATCTA ATCC].

Amplicon polymerase chain reaction (PCR) was performed to amplify the template from the DNA sample input. Briefly, each 25 μL PCR contained 12.5 ng of sample DNA as an input, 12.5 μL of 2x KAPA HiFi HotStart ReadyMix (Kapa Biosystems, Wilmington, USA) and 5 μL of 1 μM of each primer. PCRs were carried out using the following protocol: an initial denaturation step was performed at 95°C for 3 minutes followed by 25 cycles of denaturation (95°C, 30 s), annealing (55°C, 30 s) and extension (72°C, 30 sec), and a final elongation for 5 minutes at 72°C. The reaction mix was removed from the PCR product with Mag-Bind RxnPure Plus magnetic beads (Omega Biotek).

A second index PCR amplification, used to incorporate the barcodes and sequencing adapters into the final PCR product, was performed in 25 μL reactions using the same master mix conditions as described above. The cycling conditions were as follows: 95°C for 3 minutes, followed by 8 cycles of 95°C for 30 minutes, 55°C for 30 minutes and 72°C for 30 minutes. A final 5-minute elongation step was performed at 72°C. The library was checked using an Agilent 2200 TapeStation and quantified using a QuantiFluor dsDNA System (Promega). Libraries were then normalized, pooled and sequenced (2 × 300 bp paired-end read setting) on the MiSeq (Illumina, San Diego, USA) using a 600 cycle V3 standard flowcell producing approximately 100,000 paired-end 2 × 300 base reads (Omega Bioservices, Norcross, USA).

### 16S rRNA sequencing analysis

For each sample, the raw reads were filtered based on sequencing quality using Trimmomatic. The primer and adaptor sequences were removed. Sequence reads with both pair end qualities lower than 25 were truncated. The software package QIIME was used to perform the 16S rRNA analyses(2). Sequences were clustered into operational taxonomic units (OTUs) at a 97% similarity cutoff, and the relative abundance was calculated for the OTUs in each sample. All sequences were classified using a native Bayesian classifier trained against the RDP training set (version 9; http://sourceforge.net/projects/rdp-classifier/), and OTUs were assigned a classification based on which taxonomy had the majority consensus of the sequences within a given OTU. The OTUs were then aligned to the Silva database. Alpha diversity (including the Chao 1 index, the ACE index, the Shannon index, the Simpson index and the Evenness index) and the UniFrac-based principal coordinates analysis (PCoA) were performed based on the sample group information.

### The prediction of bacteria function

The relative representation of the bacteria characteristics was predicted using BugBase on the basis of six phenotype categories <u>(Ward *et al*. unpublished)</u>: Gram staining, oxygen tolerance, ability to form biofilms, mobile element content, pathogenicity, and oxidative stress tolerance. This software balances the Kyoto Encyclopedia of Genes and Genomes (KEGG) database, the Integrated Microbial Genomes (IMG4) platform and the Pathosystems Resource Integration Center (PATRIC) system to confirm the contribution of specific OTUs to a community-level phenotype(17–19). PICRUSt was used to predict the functional composition of a metagenome using marker gene data and a database of reference genomes. Functional differences among the different groups were compared using STAMP software (20, 21).

### Statistics

Analyses were performed in SPSS unless stated above. *P* < 0.05 was considered an indication of statistical significance. The differences in age and parity of patients were assessed with the use of Student’s t test. The differences in menopausal status, history of hypertension and diabetes were assessed using the chi-square test. Differences in the number of ovarian bacteria taxa were assessed with the use of the Mann-Whitney U test.

## Acknowledgments

We thank the colleagues in the Department of Gynecology of First Affiliated Hospital in Xi’an Jiatong University for their contributions to collecting samples.

## Funding

This work was supported by grants from the Fundamental Research Funds for Xi’an Jiaotong University (xjj2015093), and Major Basic Research Project of Natural Science of Shaanxi Provincial Science and Technology Department (2017ZDJC-11), the Key Research and Development Project of Shaanxi Provincial Science and Technology Department (2017ZDXM-SF-068), and Shaanxi Provincial Collaborative Technology Innovation Project (2017XT-026, 2018XT-002). The funders had no role in study design, data collection and analysis, decision to publish, or preparation of the manuscript.

## Author Contributions

Conceptualization: Qiling Li, Qing Song

Formal analysis: Qi Wang, Lu Han

Funding acquisition: Qiling Li, Qing Song

Investigation: Lanbo Zhao, Chao Sun, Xiaoqian Tuo

Methodology: Guoxing Fu, Sijia Ma, Qing Li

Resources: Yiran Wang, Dongxin Liang, Qing Wang, Miaomiao Tang

Supervision: Qiling Li

Visualization: Qing Song

Writng original draft: Qi Wang

Writing-review & editing: Lanbo Zhao, Lu Han

